# Complementary chromosome folding by transcription factors and cohesin

**DOI:** 10.1101/305359

**Authors:** M. C. F. Pereira, C. A. Brackley, D. Michieletto, C. Annunziatella, S. Bianco, A. M. Chiariello, M. Nicodemi, D. Marenduzzo

**Affiliations:** SUPA, School of Physics and Astronomy, University of Edinburgh, Peter Guthrie Tait Road, Edinburgh EH9 3FD, Edinburgh, UK; Dipartimento di Fisica, Università degli Studi di Napoli Federico II, and INFN Napoli, Complesso Universitario di Monte Sant’Angelo - 80126 Naples, Italy; Berlin Institute of Health (BIH), and Berlin Institute for Medical Systems Biology at the Max Delbruck Center for Molecular Medicine - 13125 Berlin, Germany

## Abstract

The spatial organisation of interphase chromosomes is known to affect genomic function, yet the principles behind such organisation remain elusive. Here, we first compare and then combine two well-known biophysical models, the transcription factor (TF) and loop extrusion (LE) models, and dissect their respective roles in organising the genome. Our results suggest that extrusion and transcription factors play complementary roles in folding the genome: the former are necessary to compact gene deserts or “inert chromatin” regions, the latter are sufficient to explain most of the structure found in transcriptionally active or repressed domains. Finally, we find that to reproduce interaction patterns found in HiC experiments we do not need to postulate an explicit motor activity of cohesin (or other extruding factors): a model where cohesin molecules behave as molecular slip-links sliding diffusively along chromatin works equally well.

---

Interphase chromosomal organisation is intimately linked to gene regulation and cellular integrity [1–3]. Distinct genomic architectures can be found in cells undergoing differentiation and ageing or in those affected by disease [4, 5]. Recent years have seen major developments in a number of techniques to investigate the 3-D conformation assumed by interphase [6–9] and mitotic chromosomes [10]. The most widely employed technique to date is “HiC” - a high-throughput, genome-wide version of “chromosome conformation capture” whose natural output is a map quantifying the probability of interaction between different genomic loci within a population of cells [6–9, 11]. These maps, constructed for different organisms and cell types [7], naturally lend themselves to comparison with those predicted by “bottom-up” computational models based on polymer physics principles [12–14].

Two main classes of biophysical models are currently popular in the field: the “transcription factor” (TF) model [12, 15–17] (also known as the “strings-and-binders” model [18]); and the “loop extrusion” (LE) [13, 19, 20] model. The former postulates that multivalent chromatin-binding proteins mediate chromatin-chromatin interactions, creating loops and driving 3-D folding. Examples of proteins that play key roles within this framework are transcription factors associated with active chromatin [21], as well as polycomb group [22] and HP1 [23] proteins. This model naturally explains the large-scale (micro)phase separation of the genome into active and inactive (also known as A and B) compartments [12] and the formation of nuclear bodies [24], both driven by a mechanism known as the “bridging-induced attraction” [16]. The second model posits that the SMC complex cohesin and the CCCTC-binding factor (CTCF) are the master organisers of the genome, suggesting that cohesin acts as a loop extruding factor [25] which actively creates expanding loops, but halts when it meets a bound CTCF. This model can account for the striking bias in favour of convergent CTCF loops [9] and it can also rationalise the “topologically-associated-domain” (TAD) patterns observed in HiC maps [13]. However, a motor activity has yet to been observed in experiments probing the motion of DNA-bound cohesin *in vitro* [26–28] and the convergent loop bias can also be explained by a model of diffusive loop extrusion (dLE) where cohesin slides diffusively along the chromatin rather than actively moving unidirectionally [29]. A third possibility is that the diffusive motion is enhanced by ATP consumption resulting in an active bidirectional motion.

The TF and (d)LE models each explain different aspects of genome organisation. While the TF model describes a functional level of genome organisation, intimately linked to the local transcriptional activity and chromatin state, the LE model describes a level of organisation independent of these. The reality may well be a combination of the two, in which case one would expect that disrupting either transcription factor or cohesin binding would give rise to distinct changes in chromosomal architecture. Indeed, very high resolution conformation studies of the globin loci using Capture C [17, 31] revealed completely different conformations in erythroid cells, where these genes are very active, and stem cells, where they are inactive - i.e. changes in protein binding sites result in changes in conformation. Likewise, cohesin or CTCF knock-outs result in the disruption of the observed loops and loop-domains [25, 32–34], but appear to leave the underlying chromatin states mostly unchanged [34].

In this computer simulation study, we first compare the TF and (d)LE models in terms of their ability to predict chromosome organisation. We focus our attention on a 30 Mbp section of human chromosome 7, which includes large gene deserts (regions of “inert chromatin”, where transcriptional activity is sparse and which is void of active or repressive histone modifications), as well as facultative and constitutive heterochromatin, and active regions. We find that neither the TF nor the LE model can, by itself, give a satisfactory account of the observed folding of the entire chromosome segment. The (d)LE model accurately predicts the domain pattern locally, but fails to capture larger-scale interactions. On the contrary, the TF model poorly predicts the fine detail of local interactions (especially within gene deserts/inert chromatin), but captures long-range contacts more faithfully.

A combination of the TF and (d)LE models reproduces many of the HiC features, suggesting that TFs and cohesin (or other LE factors) indeed have complementary roles in genome organisation. We show that LEs are required to create TADs within inert chromatin, while our simulations suggest that TFs are sufficient to organise active/inactive domains, where cohesin-mediated loops play a more minor role.

Intriguingly, however, a naïve superposition of the standard TF and (d)LE models still leaves some key qualitative discrepancies between simulated and HiC interaction maps; for example the simulations tend to show too high a signal for medium to long range interactions. Qualitative agreement improves when the TF model is enhanced by including a non-equilibrium “switching” mechanism [24]. This switching-TF (sTF) model encodes a dynamic level of control on TFs; it might represent post-translational modification of the proteins affecting their binding affinity to chromatin [35]. The combined model with switching TFs is also consistent with single-molecule microscopy experiments on the dynamics of active/inactive chromatin domains, chromatin loops and protein clusters [36–38], and it correctly predicts the main observations of recent knockout experiments [34].

## METHODS

Following our previous work [12, 17, 24, 29, 39–41], we employ a simulation scheme based on polymer physics: the chromatin fibre is represented as a chain of “beads” connected by springs. Details are shown schematically in Figure 1. Beads are “coloured” according to the underlying chromatin state based on histone modifications (ChlP-seq data are obtained from the ENCODE project [30], see Suppl. Methods bellow). In this way our chromatin becomes a co-polymer whose segments interact with freely diffusing beads mimicking explicit bridge-forming protein complexes. In our TF model, we consider three species of bridge proteins, representing transcription factors associated with euchromatin, HP1, and polycomb repressive complexes (PRC) respectively. Thus, during the course of a simulation, these proteins can bind and form bridges between chromatin beads bearing the associated epigenetic marks (see Fig. 1). Loop extruding factors, which might represent the cohesin complex, or a pair of cohesin rings, are represented as additional transient springs between non-adjacent beads (see Fig. 1 and Suppl. Methods bellow for more details).

**Figure 1:**
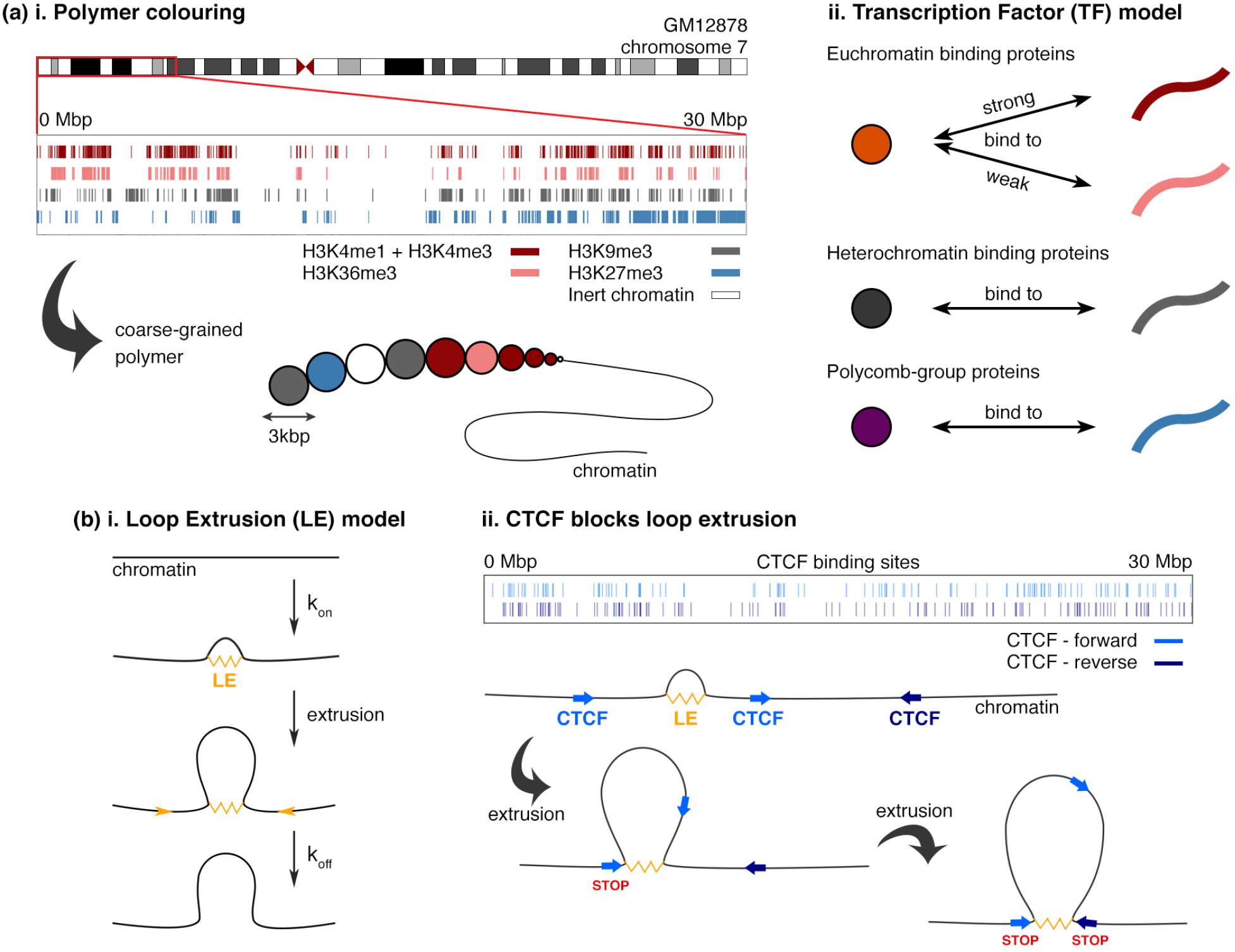
Two models for genome organization. (a) In the TF model, multivalent complexes of chromatin binding proteins organize the chromatin by forming molecular bridges between their different binding sites. In simulations, diffusing beads represent three species of TF complex and a bead-and-spring polymer represents the chromosome section. Binding sites are inferred from histone modifications, (i), and the polymer beads are “coloured” accordingly (histone modification ChIP-seq data for a human lymphoblastoid cell line GM12878 are obtained from the ENCODE project [30]). Euchromatin binding proteins (representing e.g. PolII/TF complexes) strongly bind chromatin regions with histone modifications associated with active enhancers and promoters, and weakly bind modifications associated with transcription. Two types of repressive TF bind marks associated with heterochromatin and polycomb repression respectively (e.g. representing HP1 or PRC2). (b) In the LE model, extruding factors (e.g. representing the cohesin complex) bind and unbind chromatin with rates *k_on_* and *k_off_* respectively; while bound, the factors move actively (or diffusively – see Suppl. Methods and [13] for details) along the chromatin, extruding a loop. LEs stop moving if they reach a CTCF site where the binding motif is orientated towards its direction of motion, or if they encounter another extruder. ChIP-seq data for CTCF binding in GM12878 cells, (ii), are also obtained from ENCODE (see Suppl. Methods for details on motif identification). In simulations, the chromosome is represented by a bead-and-spring polymer, and extruders are realized by adding extra springs, starting from adjacent beads and progressively moving forward in time.

## RESULTS

In this paper we focus on the first 30 Mbp of human chromosome 7 in a human lymphoblastoid cell line GM12878, for which high-resolution HiC data are available [9]. We chose this region as it contains both active and repressed regions, as well as large regions devoid of most histone modifications, which we call gene deserts or inert chromatin. Inert chromatin is AT-rich and gene-poor, so that it bears some of the signatures of heterochromatin, though it is not characterised by an enrichment of either the H3K27me3 or H3K9me3 histone modifications. Below we describe the results we obtain by applying the TF and (d)LE models, either independently or in combination.

### Neither the TF, nor the LE model alone can satisfactorily predict the observed HiC map

By applying the TF model (see Methods) we obtain the contact map shown in Figure 2(a). One way to measure how well the simulated map predicts the HiC data is to simply count the number of correctly predicted domain boundaries. From our previous work, we expect that the formation of domains which bear different epigenetic marks will be well captured by the TF model: they phase separate into distinct 3-D compartments, and clusters of like proteins form [16]. [Such clusters are visible in Figure 2(a) (see also Suppl. Movie 1) - they resemble liquid-like clusters formed by heterochromatin [23, 42] and transcription factories self-assembled within euchomatin [21].] Indeed the model does correctly capture a large fraction of boundaries in active and inactive regions (see, e.g., the 20 — 30 Mbp segment in Fig. 2(a)), as well as the pattern of longer-range interactions between segments bearing similar histone marks. These features are a natural consequence of the spatial segregation (or more precisely “microphase separation”, i.e. phase separation into domains with self-limiting size [24]) between active and inactive chromatin, which leads to A/B compartmentalisation (see Fig. S1 bellow). As well as boundaries between regions in different compartments, alternating binding and non-binding chromatin regions can also give rise to boundaries even between two adjacent active (or inactive) domains - which is why in more active regions, such as in chromosome 19, the TF model correctly predicts an even larger fraction of boundaries [12]. However, the TF model clearly fails to capture the folding of the inert chromatin regions (which is why the total fraction of correctly predicted boundaries is only ~ 36%).

**Figure 2:**
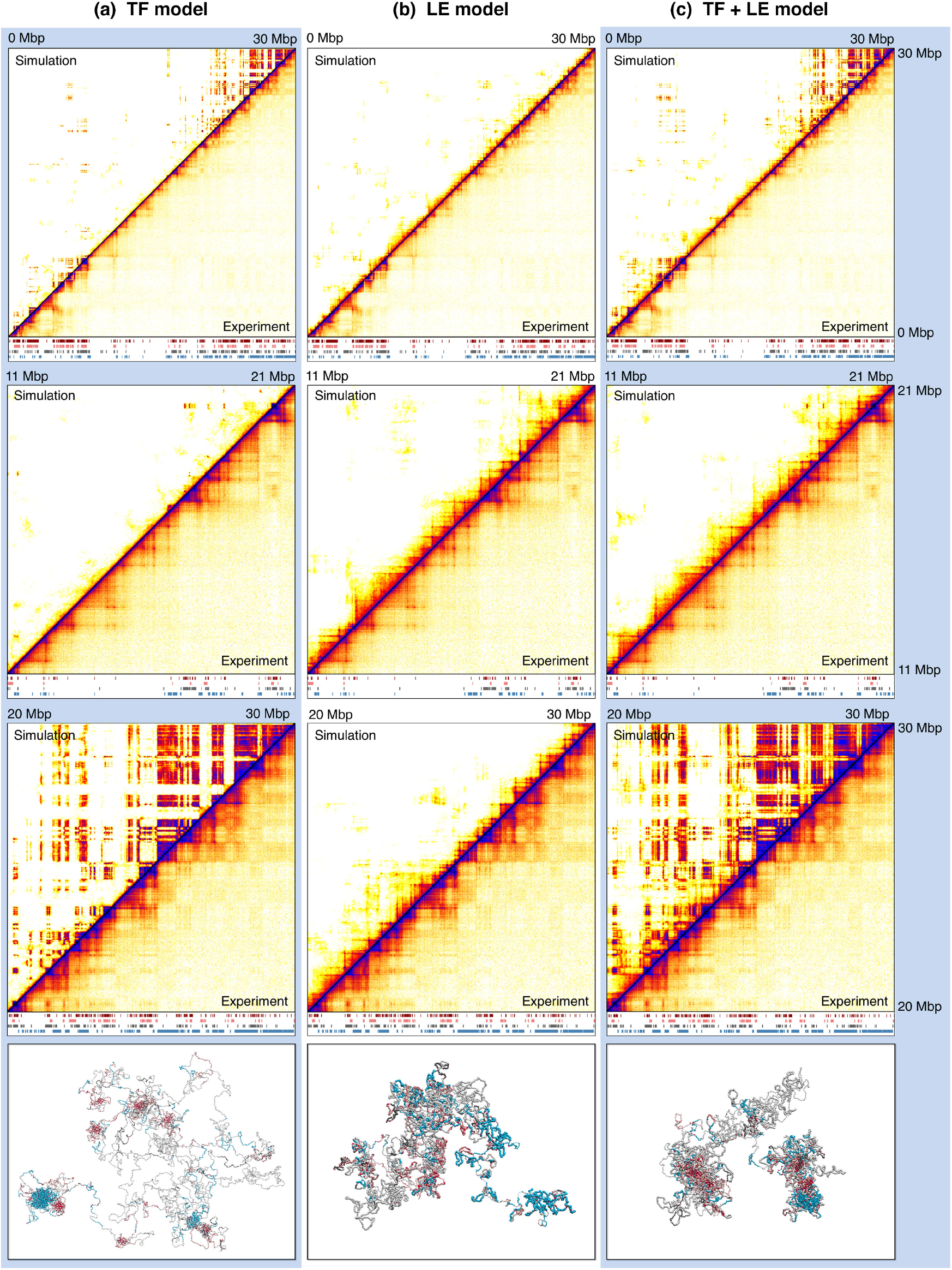
The different models and their combination predict HiC interaction maps. Plots show interaction maps for the TF model, (b) the LE model, and (c) their combination (TF+LE model). In each case the upper left triangle shows the simulation data, and the bottom right the corresponding HiC map. Top plots show the full simulated chromosome section (human chr7:1-30,000,000), with zooms on two regions shown below. Underneath each map the data used in the simulations is indicated: red, pink, grey and blue bars represent regions with H3K4me1/H3K4me3, H3K36me3, H3k9me3 and H3K27me3 respectively. TFs predict the formation of A/B compartments in regions enriched with histone modification marks, but fail to predict the folding of chromatin in inert regions. On the contrary, LEs correctly predict the formation of TADs in regions devoid of marks, but are unable to generate interactions between domains bearing the same histone modifications. The bottom panels show snapshots of the simulation for each model. For clarity TFs are not shown. The colour of the polymer segments follows the same colour scheme as their respective histone modifications, with white segments corresponding to unmodified regions. Extruded loops are highlighted by larger beads, for visualisation purposes.

Compared to the TF model, the LE model gives a better prediction of local TAD formation, especially within inert chromatin (where 82% of domain boundaries are correctly predicted), but it performs less well in capturing the higher-order organisation of active and inactive regions. Although a similar number of boundaries (83%) are correctly predicted in those regions, the contact maps obtained with the LE model distinctly lack the long-range interactions between domains, which are associated with compartmentalisation.

The LE model also clearly cannot capture enhancer-promoter interactions within domains unless there are CTCF sites in the vicinity of those regulatory elements. To highlight this, we considered a virtual 4C experiment by selecting HiC interactions for a locus corresponding to a promoter, and compared the interaction patterns predicted by the TF and LE models. The results clearly show that the LE model fails to capture the pattern qualitatively (see Fig. S2): the correlation between the virtual 4C interaction profiles for the simulated and HiC data is −0.003 for the LE models, compared to 0.27 for the TF model.

### A naïve combination of TF and LE models improves the qualitative agreement with HiC, yet some issues remain

Since each of the models captures different features of chromosome folding, one expects that a combination of the two should perform much better than either on its own. Indeed we find that the combined TF+LE model (see Methods for implementation details) yields an improvement, as now both inert and active/inactive regions are in fair qualitative agreement with HiC (Fig. 2(c)); however some discrepancies remain - these are discussed in more detail below.

An important result is that, within our simulations, extruders are not necessary for local folding within regions that display well-defined patterns of histone modification - their organisation is mainly driven by TF bridging (Fig. 2(c)). For instance, the simulated contact maps for the TF and TF+LE model in the 20 - 30 Mbp region, which is rich in active and inactive domains, are highly correlated (Pearson’s correlation *r* = 0.76). The main reason for this is that when bridges bind they tend to compact a whole stretch of chromatin, creating many more contacts compared to extruders, each of which only forms a single loop.

Notwithstanding the improved agreement with HiC, a visual inspection of the contact maps in Figure 2 reveals that there are some remaining qualitative discrepancies. Most notably, there are substantially more interdomain interactions far from the diagonal in the simulated contact maps, whereas these features are much weaker in the HiC map (see Fig. 2(c), bottom zoom, between 20 – 30 Mbp).

### A model with switchable TFs shows qualitatively better agreement with HiC and fluorescence microscopy experiments

We now consider a variation of the TF model which gives improved qualitative agreement with HiC experiments [9]. In the model discussed above, active and inactive factors interact with chromatin beads thermodynamically - i.e., there is an attractive binding interaction between the TF and respective chromatin beads. A TF can bind chromatin when it diffuses into contact, and then unbinds due to the thermal motion in the system. The residence time depends on the interaction strength (see Suppl. Methods) and it is strongly modulated by emergent behaviour such as the bridging-induced attraction [16]. More specifically, once a multivalent factor reaches a configuration where it can form multiple chromatin interactions, it remains bound for a time which increases exponentially with number of interactions. This is because unbinding requires climbing over a potential energy barrier whose height increases linearly with the number of interactions. Within our baseline model, typical residence times can encompass the total simulation time, thus the model fails to capture the rapid turn-over of TFs observed *in vivo* (typically of the order of minutes - see below and Refs. [37, 43]).

Many TFs and other proteins which are relevant to our modelling are observed in stable foci which also exhibit rapid protein turnover. These two features are difficult to reconcile, but possible explanations are that there is ongoing post-translational modification which affects binding affinities (e.g. phosphorylation [35, 44]), that there is active protein degradation, or, in the case of PolII, that transcription-termination signals lead to unbinding [1]. A generic way to model these non-equilibrium processes is to consider TFs that switch between an “on” (binding) and an “off” (non-binding) state at rate *k_switch_* (see Fig. 3(a)). We have recently shown [24] that this switching-TF (sTF) model gives rise to the formation of dynamic protein clusters, reminiscent of nuclear bodies [45], and can affect chromatin interaction patterns.

**Figure 3:**
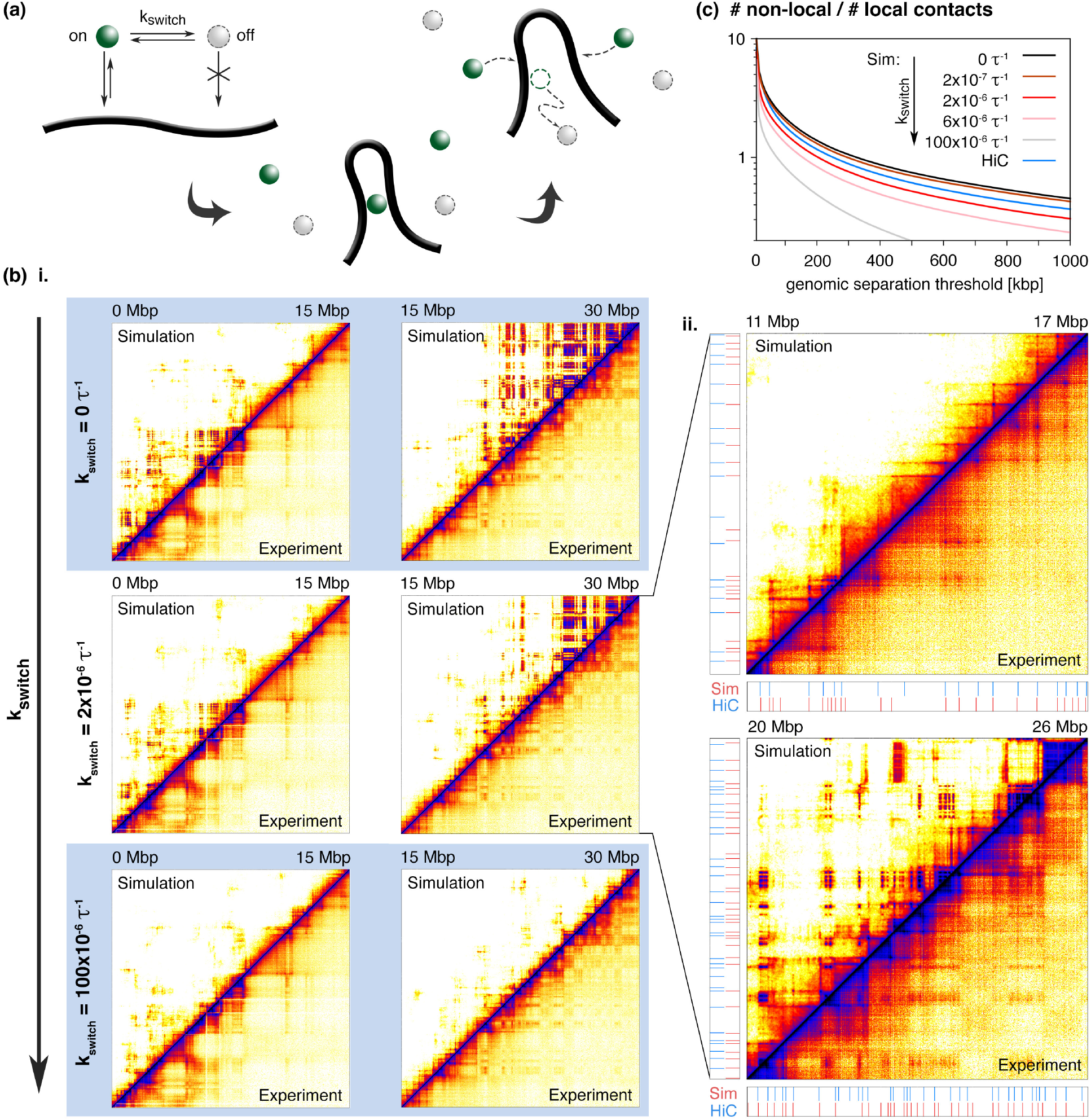
TF switching improves agreement with HiC data. (a) TFs switch between an “on” (binding) and an “off” (non-binding) state at rate *k*_switch_. This leads to the formation of dynamic, rather than static, protein clusters. (b) i. interaction maps comparing sTF+LE simulations (top left triangles) with HiC data (bottom right). From top to bottom the switching rate *k*_switch_ is increased from 0 to 100 × 10^−6^ *τ*^−1^ (where *τ* is the simulation time unit - see Suppl. Methods for how this can be mapped to a real time). Left and right maps show different regions of the simulated chromosome section. Higher switching rates yield less inter-domain interactions. ii. Zooms on different regions of the simulated section with *k*_switch_ = 2 × 10^−6^ *τ*^−1^ (corresponding to 40 × 10^−6^s^−1^). Bellow and to the left of the maps the positions of domain boundaries are indicated - see Suppl. Methods for details on boundary detection. The sTF+LE correctly predicts 84% of the HiC domain boundaries. (c) Plot showing the ratio of non-local to local interactions for HiC and simulation maps, for varying *k*_switch_ values. Instead of fixing a threshold for locality, we plot the ratio as a function of the threshold for each case. The TF+LE model without switching predicts too high a ratio of non-local to local interactions, when comparing with the HiC curve.

Figure 3(b) shows the qualitative effect of TF switching on the contact maps for different values of the switching rate *k*_switch_. The most striking difference between the combined TF+LE models with and without switching is that switching markedly attenuates long-range inter-domain, but not intra-domain, interactions: active domains which are far apart along the genome are less likely to interact. This reduces the intensity of the off-diagonal features in our predicted contact maps, rendering them qualitatively more similar to the HiC. This is also shown by the decrease in the ratio of non-local to local contacts with an increasing *k_switch_* (see Fig. 3(c)): the TF+LE model with switching is needed to predict the correct balance between long - and short-range interactions.

The sTF+LE model also conforms much better with observations from live-cell fluorescence microscopy experiments which probe dynamical information inaccessible to HiC. In the absence of switching, the TF dynamics in our model is slow and glassy, whereas it is much more rapid with switching (Fig. 4, and Suppl. Movies 1, 2). Whilst high-throughput experiments showing the dynamics of chromatin interactions over time are not yet possible, the more dynamical picture emergent from the switching model is consistent with fluorescence recovery after photo-bleaching (FRAP, see above and [24]) and single-molecule imaging experiments, which suggest that TF binding is short-lived and lasts for not more than minutes [37, 43]. The differences are clear if we examine the trajectories of individual TFs (Fig. 4(a)): without switching, a TF diffuses until it joins a cluster (of like proteins and binding sites), where it tends to stay for the remainder of the simulations; with switching a TF joins a cluster for a short time, then undergoes a period of free diffusion, before joining another cluster. This hopping between clusters manifests as a long tail in the distribution of mean squared displacement for a given time interval (Fig. 4(b)). Supplementary Movies 1 and 2 also show that the macroscopic dynamics of liquid-like domains formed by non-switching and switching proteins are profoundly different: in the latter case, we observe many more events corresponding to clusters splitting and reforming, and also the clusters are smaller.

**Figure 4:**
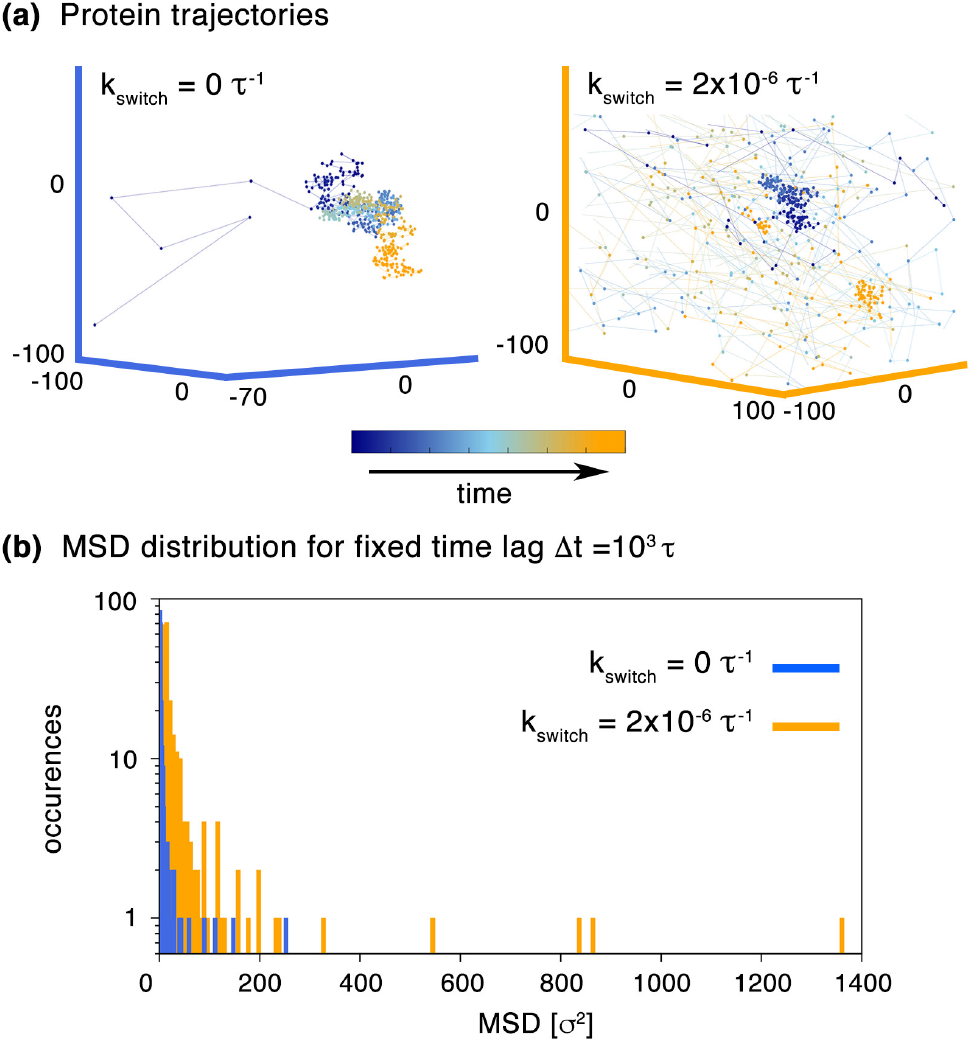
Dynamics of protein structures in the sTF model. (a) Trajectories in 3-D for a single protein without (left) and with (right) switching. Without switching, the TF quickly gets “stuck” in a cluster, after which point its dynamics are constrained. With switching, the TF stays in a cluster for a short time, before unbinding and diffusing freely, and later joining a different cluster. (b) Protein dynamics are characterized by the mean squared displacement (MSD) of TFs for a fixed time interval of 10^3^*τ*; the distribution of the MSD is shown, for two values of *K*_switch_.

### A quantitative analysis confirms that the sTF+LE model gives the best agreement with HiC

We now turn to a more quantitative analysis of the agreement between HiC and the set of models considered. For this we consider a series of parameters.

First, we compare the model performance in domain boundary generation. For the sTF+LE model, we find that across the whole simulated region, ~ 84% of boundaries are correctly predicted (see Fig. 5(c)). This performance is similar to that of the LE and TF+LE models (~ 83% and ~ 82% respectively), but substantially better than the TF model (~ 36%). Previous studies with only TFs [12], or only extruders [13] found similarly high values, however neither focused on chromosome regions containing both inert and active/inactive regions, as we have done here.

**Figure 5:**
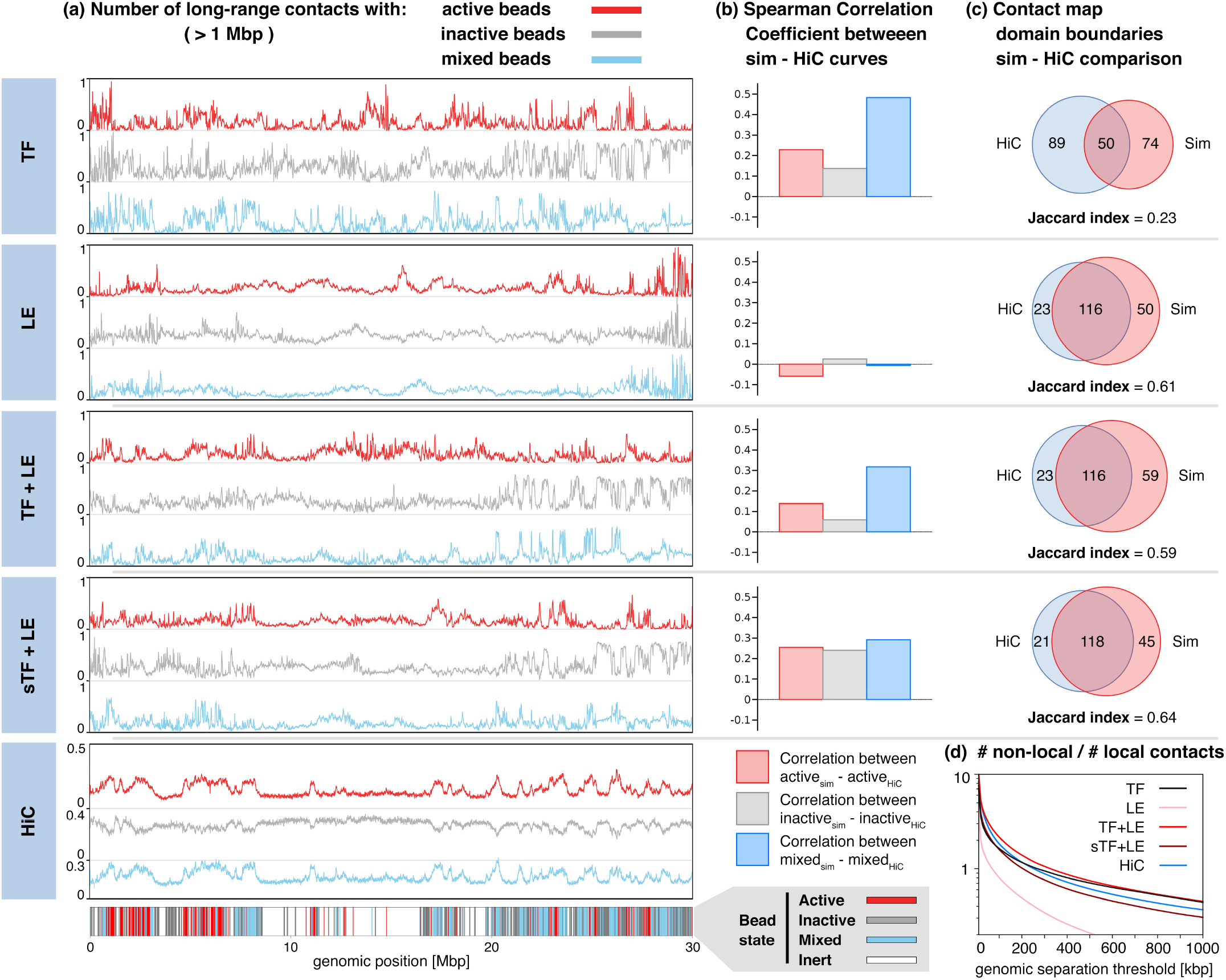
A quantitative comparison of the various models. (a) Plots show, for each chromatin bead across the simulated region, the level of interaction with active (red), inactive (grey) or mixed (blue) beads. For each bead the number of interactions with each type is normalised so that the total number of interactions sums to 1. The labelling of each bead is indicated below the plots. (b) Bar plot showing the Spearman rank correlation coefficient for comparison of each curve in (a) with the corresponding HiC data. The LE model yields long-range interaction patterns that essentially do not correlate with HiC. In general, models with TFs give a good correlation with the HiC profiles. (c) Venn diagrams showing the overlap between called domain boundaries in the HiC and simulated contact maps. The Jaccard index gives a measure of the degree of overlap, ranging from 0 (no boundaries match) to 1 (exact match between simulation and data) – see Suppl. Methods for more details. Since boundaries are related with the fine domain structure near the interaction map’s diagonal, models with LEs give in general a good prediction. (d) Plot showing the ratio of non-local to local interactions for HiC and simulation maps, for the different models, as a function of the threshold. The sTF+LE model gives the best prediction of the HiC curve (see also Fig. 3(c)).

Second, we consider a measure of the long range active-active, inactive-inactive, and active-inactive interactions to assess how well each model captures compartmentalisation and formation of promoter-enhancer hubs [21]. To do this we label each chromatin bead as active or inactive according to whether it binds to active or inactive TFs (which is in turn based on histone modification data); we label beads which can bind to both as “mixed”, and which bind neither as “inert”. Figure 5(a) shows the fraction of active, inactive and mixed beads which each chromatin bead interacts with, for each of the models. In the HiC data there is an enrichment of active-active and inactive-inactive contacts, which is associated with compartmentalisation. This is captured by models with TFs (where there is a high correlation with the HiC - see Fig. 5(b)), but not by the LE model (which shows essentially no correlation). A Spearman correlation test shows that the combined model with switching performs better than the combined model without switching (although it is not significantly better than the TF only model). A similar conclusion is reached by inspection of virtual 4C interaction profiles for promoters/enhancers (see example in Fig. S2).

Third, we assess the relative balance between local and non-local contacts in the various models and in experiments (Fig. 5(d)). This further supports the idea that the TF and LE model separately cannot fully account for HiC data, and also shows that to capture the right decay of the non-local to local fraction a model with switching is required (see also Fig. 3(c)).

### An *active* extrusion mechanism is not necessary to create domains and CTCF loops

Having noted that CTCF and extrusion appear to be fundamental to the creation of domain boundaries, especially in inert chromatin, we now ask whether *active* extrusion (where LEs move unidirectionally due to some motor effect as in Refs. [13, 20]) is necessarily required, or whether *diffusive* extruders (dLE) behave similarly. This is currently a relevant question as single molecule experiments on cohesin loaded onto DNA or reconstituted chromatin [26–28] have not yet found evidence of a direct motor activity. [Indirect motor activity, e.g. by a transcribing polymerase, is a distinct and plausible alternative, and has been suggested on the basis of simulations [46]; however it is difficult to find direct evidence for this *in vivo*.]

Simulating large chromosome regions with a dLE model in 3-D requires using either infeasibly long simulation times, or using substantially coarser resolution. Therefore we first studied a 1-D model of dLE (see Suppl. Methods). CTCF sites were positioned as in the 3-D simulations with active LEs, and, like before, we assume that the diffusing LEs interact strongly and directionally with CTCFs (see Methods for more details). Contact maps can be computed within this 1-D model by assuming a HiC interaction between the position of each pair of monomers in a diffusing loop extruder (cohesin dimer). The resulting interaction maps are plotted in Figure 6(a), together with maps from active LE simulations, computed in the same “1-D fashion” (see Suppl. Methods). Results show that dLE is essentially indistinguishable from active LE, both visually and quantitatively: 83% of the HiC domain boundaries were correctly predicted by the 1-D dLE model.

**Figure 6:**
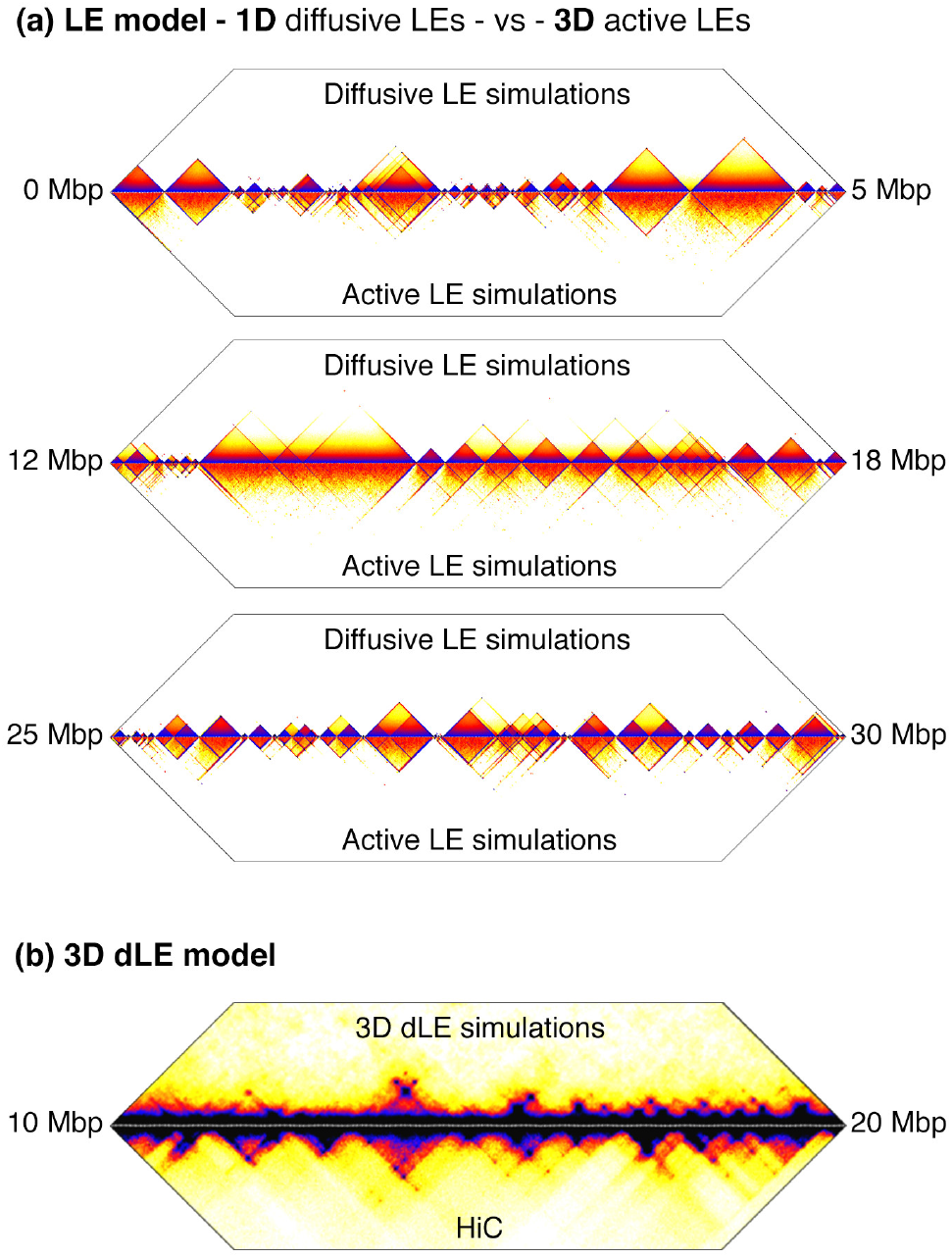
Diffusive LEs give similar predictions to the active LE model. (a) Plots comparing simulated HiC maps generated by the active (bottom) and diffusive (top) LE model. The active LE results are from the 3-D simulations as shown in previous plots, whereas the dLE results are from a 1-D model (see Suppl. Methods). In order to make a fair comparison, here the interaction map for the 3-D simulations was calculated in a “1-D fashion”, i. e., two chromatin beads are considered to be in contact if they are bound by a LE. Three different regions of the simulated chromosome section are shown. Note that due to the way the interactions are defined it is not possible to generate long-range contacts in the 1-D model. (b) Plot comparing the simulated HiC map generated by the 3-D diffusive LE model (top) and the *in-situ* HiC map from experiments (bottom). (see Suppl. Methods for details on the 3-D dLE model)

We also performed 3-D simulations of dLE in the region between 10 and 20 Mbp, with 25 kbp resolution per chromatin bead - this lower resolution allows sufficiently long simulations for the dLE maps to reach steady state. Results confirm that dLE does reproduce most of the boundaries and peaks shown in HiC (see Fig. 6(b); for this case, simulations also include a non-specific attraction between all beads to qualitatively account for the effect of macro-molecular crowding [20]).

### The combined sTF-LE model correctly predicts the effects of various protein knock-outs

We next use our combined sTF+LE model to simulate the effect of cohesin removal and targeted CTCF degradation, which were both recently explored experimentally. We find that simulations qualitatively reproduce the experimental observations.

Cohesin removal in the simulations leads to loss of folding in inert chromatin regions, leaving little structure in the contact maps (Fig. 7(a)i). This mirrors observations from experiments that knocked out NIPBL, which is required for cohesin loading in mammalian cells [25, 32]. On the other hand, domains organised by active and inactive switching factors are only subtly affected in our model. This is qualitatively consistent with the results of Ref. [25], which found some residual structure in active/inactive compartments (but not inert ones) following cohesin removal in mouse liver cells (see Suppl. Fig. 5 in Ref. [25]). More specific to our work, we show in Figure S3 HiC data for an active region in a similar chromosome region as considered here: it can be seen that some peaks and the overall contact pattern remain in the NIPBL knock-out. Like in the experiments, the simulated interaction map reveals stronger compartmentalisation upon cohesin removal, with a decrease in the number of interactions between domains with different epigenetic marks, and an enhancement of the interactions between like domains (see Fig. S4(a)).

**Figure 7:**
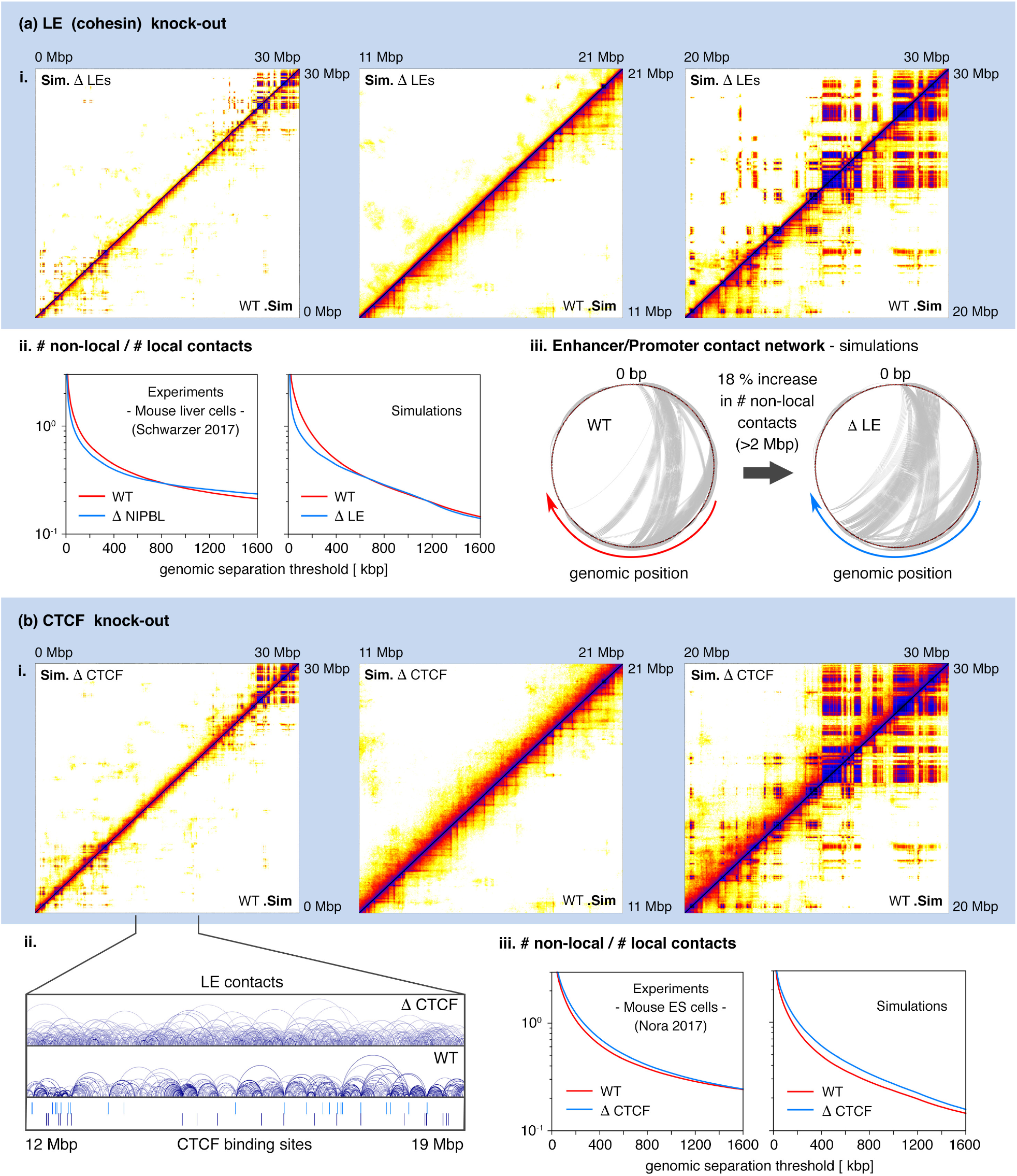
The sTF+LE model predicts the effect of cohesin and CTCF knock-out. (a) A cohesin knock-out simulation is performed by removing LEs. i. Interation maps where the knock-out (top left triangle) is compared with a wild type (WT) simulation (bottom right). The leftmost map shows the entire simulated region, with two zooms shown to the right. ii. Plots showing the ratio of non-local to local interactions as a function of the threshold for the KO and WT cases: on the left results are from experiments in mouse liver cells and on the right from simulations. iii. Circos plots showing the interaction network between chromatin beads corresponding to regions with H3K4me1 and H3k4me3 marks, for the simulated KO and WT cases. In the outer circumference chromatin beads are displayed ordered by their genomic position in the clock-wise direction. Interactions are drawn as lines connecting interacting beads - only interactions with probability > 0.1 are displayed for better visualisation. (b) A CTCF knock-out simulation is performed by omitting CTCF binding sites. LEs move until they unbind, or are blocked by other LEs. i. Similar interaction maps to the above for the entire simulated region (left) and two zooms. ii. 1-D representation of LE contacts for the simulated CTCF KO and WT cases. The arc ends correspond to chromatin beads bound by a LE, and the arc length is proportional to the beads’ genomic separation. CTCF binding sites are indicated bellow. iii. Similar plots showing the ratio of non-local to local interactions for the CTCF KO and WT cases: on the left, results are from experiments in mouse embryonic stem cells and on the right from simulations.

To better access the qualitative agreement with experiments, we extracted from the interaction maps the ratio of non-local to local interactions as a function of the genomic separation threshold for “locality”. Figure. 7(a)ii shows the plots comparing KO and WT cases, for the simulated and HiC maps, obtained from mouse liver cell experiments [25]. There are two distinct regimes: for thresholds below the TAD range (~ 700 — 800 kbp) there is a loss in non-local interactions upon cohesin removal, and above the TAD range there is a loss in local interactions. Our model captures these features up to a threshold ~ 1100 kbp. Above that the simulation predictions deviate from the experimental observations - our WT model yields a higher ratio of non-local to local interactions. This is due to the choice of low concentrations in our model (which avoids non-physical confinement effects), which allows the polymer to change its conformation faster, meaning that loop extrusion will in fact favour a more compact structure and therefore more non-local interactions (see simulation snapshots in Fig. 2).

Experiments also reported the formation of superenhancer hubs following cohesin removal [32]. Superenhancers are genomic regions containing a high linear density of enhancer elements and high levels of the associated H3K27ac histone modification. Interactions between superenhancers - including interchromosomal interactions - were found to increase after cohesin removal, and examination of HiC ligation events revealed a higher instance of triplets of these loci appearing together [47] (i.e. three of these loci were in close proximity at the same time). To assess whether extruders qualitatively affect the network of active chromatin contacts in our simulations, we show in Figure 7(a)iii circos diagrams for enhancer/promoter chromatin beads only, for the KO and WT simulations. The chromatin beads are ordered according to their genomic position along the outer circumference in the clock-wise direction. Upon cohesin KO there is an 18% increase in the number of non-local interactions (genomic separation > 2 Mbp). This is further supported by analysing clusters of TFs binding active euchromatin, formed through the bridging-induced attraction. These indeed involve more non-local interactions between binding sites after LE removal: the mean genomic separation of chromatin beads associated with such clusters raises by over 10% from 837 kbp to 948 kbp. It is therefore tempting to associate these active protein clusters with the superenhancer hubs found experimentally. Our simulations also show that cohesin loss results in a minor decrease in the sizes of TF clusters (however the change is not statistically significant according to a Kolmogorov-Smirnov test).

CTCF removal leads to a loss of “hot-spots” in the contact map, which in wild-type nuclei correspond to convergent CTCF loops (Fig. 7(b)i). Domains and boundaries become much less well-defined within the inert chromatin region, but are relatively unaffected elsewhere (see Venn diagrams for identified boundaries in Fig. S4(b)). The spatial distribution of cohesin on the chromosomes is also strongly affected (see Fig. 7(b)ii). These findings are in agreement with experiments knocking out CTCF [33, 34] in mouseembryonic stem cells. In the wild-type simulations, cohesin localises mostly at CTCF sites, consistent with ChIP-seq data [48]. In the CTCF knock-out simulations, cohesin is distributed uniformly across the chromosome segment. In experiments, cohesin instead accumulates at transcription start-sites upon CTCF loss. One possible reason for this discrepancy is that cohesin might have preferred loading sites on the chromatin (some preferential binding of NIPBL, required for loading, has been observed at transcription start sites [48]). We have previously shown that including preferred loading sites in simulations would, in the absence of CTCF, leads to an enrichment of cohesin at those sites [29].

We also compared the ratio of non-local to local interactions as a function of the genomic separation threshold for the KO and WT cases (see Fig. 7(b)iii), for the simu-lated and HiC maps, obtained from mouse embryonic stem cell experiments [34]. CTCF removal has a minor effect on these ratios, slightly favouring more non-local contacts. Our simulations qualitatively agree with the experimental observations for all analysed contact separation threshold values.

## DISCUSSION AND CONCLUSIONS

In this work we have studied chromosome folding by using a combination of two popular and successful models for mammalian genome organisation: the transcription factor [12, 15, 16, 18] and loop extrusion [13] models. The TF model is motivated by the abundance of multivalent architectural chromatin-binding proteins or complexes (e.g., HP1, PRC1, TF/PolII complexes etc.), which are known to form loops within the genome, and organise it into active and inactive regions. The TF model naturally explains the observations of transcription factories [21] and nuclear bodies [24] as multivalent TFs generically cluster through the bridging-induced attraction [16]. The LE model is motivated by the evidence that cohesin mediates chromatin looping between convergent CTCF sites in the genomes of mammals [9].

Our simulation results suggest that TFs and cohesin play complementary roles in genome organisation. On the one hand, cohesin is necessary to organise and compact regions of inert chromatin (gene deserts) where depletion of most histone marks is consistent with minimal TF binding. Accordingly, cohesin is required to account for many of the TAD boundaries in the region of human chromosome 7 we focussed on here (which contains a large gene desert). On the other hand, activating and repressive TF factors are sufficient to organise active and repressed regions respectively, as knocking out extrusion leaves largely similar contact patterns (Fig. 7).

Importantly, we find that an active mechanism for extrusion is not the only model which can generate TADs within inert chromatin: a similar number of HiC boundaries are correctly predicted by a diffusive LE model where cohesin slides along chromatin with no preferred direction (Fig. 6). This conclusion is robust, and applies to different genomic regions, for instance we analysed the folding of the segment between 20.3 and 22.6 Mbp in chromosome 4, which was considered in Ref. [20]: results (see Fig. S5) confirm that diffusive and active LE give very similar contact patterns. That the diffusive loop extrusion model works well for TAD formation is of interest since to date there is no direct evidence of unidirectional motion of cohesin on chromatin [26–28].

We found the best concordance between simulations and the available experimental evidence for a model which includes a biochemical “switching” reaction for TFs. This on⟷off switching drives the system away from thermodynamic equilibrium, and allows TFs both to bind strongly, and yet be able to dissociate frequently. The switching model gives a better prediction of long-range contacts, which would otherwise decay too slowly. More importantly, switching is necessary to reconcile simulations with fluorescence microscopy experiments which measure fast dynamics for both transcription factors [37, 43, 49, 50] and other protein clusters [24].

Our combined sTF+LE model reproduces qualitatively the effect of recent knock-out experiments. Cohesin degradation leads to unfolding and the disappearance of domain boundaries in inert chromatin regions, but results in smaller changes within active/inactive chromatin [25]. CTCF knock-out also mainly affects inert chromatin regions, and homogenises the distribution of cohesin along the chromatin fibre [33, 34].

We note that a limitation of the current work is that its methodology relies on previous knowledge of the TFs responsible for folding. A recent approach [51], has introduced a possible way to circumvent this problem, by using polymer physics and machine learning to infer the optimal, minimal number and type of TFs required to reproduce the HiC matrix within a given accuracy. Unlike the current work, though, this approach requires the HiC data as an input.

We also highlight here a recent simulation work [52], where the loop extrusion model was combined with a block copolymer model [53], which postulates a weak direct attractive interaction between all inactive regions (B compartments). Whilst this related work also found that both components of the model are required to get good agreement with HiC data, it was suggested there that extrusion may compete against compartmentalisation - e.g., if a convergent CTCF loop spans domains belonging to different compartments. This interference mechanism is appealing because it is consistent with the observation that cohesin or CTCF removal leads to an enhancement of non-local A/B compartmentalisation [25]. In the present work we did not find evidence of significant competition between chromatin-state and cohesin-mediated folding at a local level. For example, there is little difference between the TF and TF+LE models in the 20-30 Mbp region - the LEs do not interfere with the ability of TFs to organize active/repressed regions. Similarly there is no significant change in the LE loop length distribution between the LE and TF+LE models in either inert or active/repressed regions - the TFs do not interfere with extrusion. Our simulations are still fully consistent with experimental results, and the TFs and LEs do have an effect on each other with respect to longer-ranged interactions. For instance, experiments showed that cohesin loss leads to the formation of hubs of superenhancers involving very long-range contacts [32], which is associated with the increase in compartmentalisation. This result sits well within our model, as we find that protein-mediated interactions between active chromatin beads associated with promoter or enhancers become longer-range in the LE knockout.

In summary, our results suggest that these transcription factors and cohesin complexes provide two complementary mechanisms for chromosome organization, and that they are more or less important in different regions of the genome. The question of how this “division of labour” is functionally relevant remains open: we speculate that cohesin-mediated folding of inert chromatin may be useful to facilitate the transition to mitosis, where (condensin-associated) loops are likely much more abundant. We also note that our work focuses on a single cell type during interphase, where histone modification patterns are already established. We do not consider, instead, how particular patterns of chromatin state are set up during differentiation, or re-established on exit from mitosis [40, 41]. It remains possible that LEs and TFs may have a more complex relationship in such situations, when the underlying epigenetic landscape is dynamic, and we hope to address this issue in the future.

## Acknowledgements

This work was supported by ERC (CoG 648050, THREEDCELLPHYSICS). MCFP acknowlegdes studentship funding from EPSRC under grant no. EP/L015110/1. This work was also supported by grants from the NIH ID 1U54DK107977-01, by CINECA ISCRA Grant HP10CYFPS5 and HP10CRTY8P, computer resources at INFN and Scope at the University of Naples (M.N.), and by the Einstein BIH Fellowship Award to M.N.

## Author contributions

MCFP, MN, DMa designed the research project. MCFP, CAB, DMi, CA, SB, AMC developed the in-house codes. CAB retrieved and treated the experimental data. MCFP, CA, SB, AMC performed simulations. MCFP, CAB, DMi carried out data analysis. MCFP, CAB, DMi, CA, MN and DMa wrote the manuscript.

## SUPPLEMENTARY METHODS

### Generic Polymer Model for Chromatin

We model the first 30 Mbp of chromosome 7 of the human lymphoblastoid cell line GM12878 in the presence of transcription factors and/or chromatin extrusion proteins (cohesin) (see Fig. 1). The chromosome segment is modelled at 3 kbp resolution as a linear self-avoiding polymer composed of beads connected by spring bonds. Each bead, with diameter *σ* = 30 nm, corresponds to 3 kbp of DNA, hence the polymer is 10,000 beads long. The interaction between beads is characterised by a potential with three contributions. First, every two consecutive beads are connected by a finitely extensible non-linear elastic (FENE) spring given by the potential

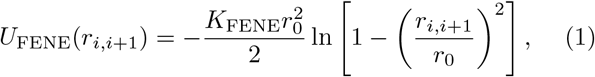

where *r_i,i+1_* is the distance between the *i*-th bead and its nearest neighbour along the chain, *r_0_* = 1.6*σ* is the maximal extent of the bond and *K*_FENE_ = 30*k*_*B*_*T/σ*^2^ is the bond energy (where *k_B_T* is the thermal energy, with *k_B_* the Boltzmann constant and *T* the temperature). Second, there is an excluded volume interaction between all beads, that prevents bead overlapping and chain crossing, given by the Weeks-Chandler-Andersen potential

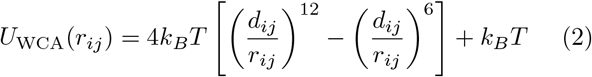

for *r_ij_ < 2^1/6^dij*, and *U*_wca_(r_ij_) = 0 otherwise. Here *r_ij_* is the distance between the *i*th and *j*th beads and *d_ij_* is the mean of the diameters of the two interacting beads, i.e., *d_i_j* = *σ*. Under this choice of FENE and WCA potentials, the bond length is approximately equal to *σ*. Third, the polymer bending rigidity is set by a Kratky-Porod potential for every three adjacent beads

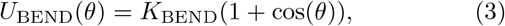

where *θ* is the angle between the three consecutive beads and *K*_BEND_ the bending energy. The polymer flexibility is set by the value of *K*_BEND_ since this determines the persistence length *l_p_* (in units of *σ*): *l_p_* = *K*_BEND_/*k_B_T*. Chromatin can be viewed as a long flexible polymer [54], therefore we use the persistence length *lp* = 3*σ* =90 nm.

### A co-polymer represents chromatin regions having different modification states

The polymer beads are “coloured” according to their underlying epigenetic state based on histone modifications. ChIP-seq data for H3K4me1, H3K4me3, H3K36me3, H3K9me3 and H3K27me3 modifications in GM12878 cells were obtained from the ENCODE project [30] (http://www.encodeproject.org [55]). We gratefully acknowledge the ENCODE project and the Bernstein Lab at the Broad Institute for generating these data (GEO:GSE29611). Modified regions (shown in Fig. 1(a) and Fig.S1) were identified either using the broad peak feature of the MACS2 peak calling software [56] (H3K4me1 and H3K4me3), or by peak calling with the Epic Peaks software (SICER) [57] (H3K36me3, H3K9me3 and H3K27me3; this algorithm is more suited to finding extended regions). Polymer beads were then annotated as active (enhancers or promoters), transcribed, heterochromatic or polycomb depending on their overlap with the called peaks (it is possible that some beads overlap multiple peaks, so these can have multiple annotations). Regions of the genome that are not enriched in any particular mark are defined as “inert”, and left unmarked.

### Transcription Factors are Modelled as Multivalent Chromatin-Binding Beads

Transcription factors are modelled, for simplicity, as spheres of diameter *σ*_prot_ = *σ*. The interaction between TFs is purely repulsive and described by the WCA potential in Eq. (2). There are three types of TFs (see Fig. 1(b)): (i) euchromatin-binding, that bind to promoters, enhancers and transcriptionally active regions; (ii) heterochromatin-binding; and (iii) polycomb-group proteins. We consider a total number of ∼ 1400 proteins [(i)∼250, (ii)∼300, (iii)-∼850], meaning that for a cubic simulation box of size *L* = 220*σ* the system’s particle concentration is ~0.1%. TFs bind to their cognate chromatin beads through an attractive interaction set by the Lennard-Jones (LJ) potential

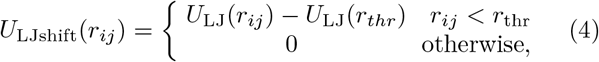

where

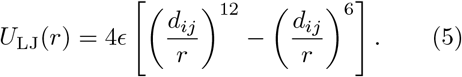

The parameter *ϵ* controls the magnitude of the proteinDNA interaction, *d_ij_* = *σ* and *r*_thr_ = 1.8*σ* is the range of the interaction. Euchromatin-binding TFs interact strongly, *ϵ* = *5.5k_B_T*, with promoters and enhancers and weakly, *ϵ* = *2k_B_T*, with transcriptionally active regions. Heterochromatin-binding TFs interact with moderate strength, *ϵ* = 3*k_B_T* with heterochromatin and interact weakly, *ϵ* = *2k_B_T*, with polycomb regions. Finally, polycomb-group proteins interact with moderate strength, *ϵ* = 3*k_B_T*, with polycomb regions and interact weakly, *ϵ* = 2*k_B_T*, with heterochromatin. The interaction between TFs and non-cognate chromatin beads is purely repulsive and given by the WCA potential in Eq. (2).

### Loop Extruders are modelled as transient bonds between chromatin segments

Loop extruders are modelled, for simplicity, as harmonic bonds that bring together two chromatin beads (see Fig. 1(c)). The bond is described by the potential

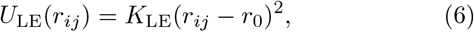

where *r_0_* = *σ* is the equilibrium bond length and *K*_LE_ = *6k_B_T* the bond strength. LE binding and unbinding is modelled by creating and deleting LE bonds (see Fig. 1(c)). Bonds are initially between *i* and *i* + 2 chromatin beads, and a loop is extruded by “moving” the ends of the bond along the polymer, i.e., by deleting the bond and creating a new one that binds the next pair of chromatin beads. We start by considering active extrusion, where the ends of the bond move away from each other along the polymer, i.e., the bond moves to the next pair of chromatin beads that corresponds to a higher genomic distance. We also perform 1-D and 3-D studies of “diffusive” extrusion (dLE), where the ends of the bond move randomly in either direction along the polymer (see details below).

An extruder comes to a halt either if it collides with another extruder or if it reaches a correctly oriented CTCF site [13]. Since the CTCF consensus binding motif is non-palindromic, it has an orientation on the DNA. A pair of CTCF sites can therefore have one of four arrangements: the two motifs could point towards each other, away from each other, or both point in the same direction. We model extruders as sensitive to this motif: if a bond end reaches a CTCF oriented in the opposite direction to its motion, then it stops, otherwise it keeps on extruding. Therefore extruders end up bringing together CTCFs with convergent motifs, in line with experimental observations [9, 20].

An extrusion step occurs every 250*τ*, where *τ* is the characteristic simulation time unit (see below for comparison with real-time), meaning that the base line extrusion rate is 2 beads/250*τ* (though see below). The number of extruders bound to the polymer is kept at a constant value of 200: a new LE binds every time one unbinds. Extruders bind at randomly chosen positions along the simulated chromatin section and unbind with a rate *k_off_* = 0.0167*τ*^−1^. In the case of an extruder which brings together a pair CTCF sites with convergent motifs, we reduce *k_off_* by a factor of 10: *k_off.conv_*CTCFs = 0.00167*τ*^−1^.

A popular candidate for the loop extruding factor is the cohesin complex [13, 20], and it is thought that a cohesin ring, or a connected pair of cohesin rings in a “hand-cuff” conformation, encircle two DNA segments, bringing them together. As such, the complex would have a finite volume and interact sterically with the surrounding chromatin and proteins. Since within our simplistic model, extruders do not occupy a finite volume (they are just transient bonds), we explicitly account for their steric hindrance by imposing a threshold on the maximum length between two beads connected by a LE bond. Practically, if the bond becomes longer than *r_ij_ > 4σ*, it is explicitly removed. This allows us to partially account for the inability of cohesin to move through a dense cluster of chromatin-bound to proteins, and constitutes a weak coupling between LEs and the 3D conformation of the polymer and proteins. Note that previous work on extrusion [13, 20, 52] does not include this coupling.

### CTCF ChIP-seq data and motif identification

ChIP-seq data for CTCF binding in GM12878 cells were obtained from the ENCODE project [30] (http://www.encodeproject.org [55]). We gratefully acknowledge the ENCODE project and the Stamatoyannopoulos Lab at the University of Washington for generating these data (GE0:GSE30263). Peaks were called using the MACS2 peak calling software [56]; the genome sequence under these peaks was then searched for CTCF binding sites. The STORM software [58] was used to identify binding motifs based on the CTCF consensus motif reported in Ref. [59]; the orientation of the peak was taken to be that of the motif scoring highest against the consensus. To account for the fact that there is a finite probability for CTCF binding at the identified sites, a binding probability was assigned to each according to the called peak height. To model cell-to-cell variability in CTCF occupancy, in each repeat simulation we populate a subset of the CTCF sites based on these probabilities. The subset of site positions were then overlayed onto our 3 kbp/bead polymer; if more than one CTCF was present in a given bead we assigned the orientation appearing most often.

### Simulating switching TFs

Following our previous work [24] we model post-translational modification of TFs by allowing them to switch between an “on” and an “off” state. While in the “on” state, TFs bind chromatin through a Lennard-Jones potential as in Eq. (4) above; when in the “off” state the interaction reverts to the non-attractive WCA potential (Eq. (2)). Importantly, the transition between these two states is controlled by an additional parameter (the switching rate *k*_switch_) that is not related to the equilibrium transcription factor unbinding time, — *τ* exp {*ϵ/k_B_T*}. Thus, the binding affinity *ϵ* and the switching time 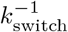 can be tuned separately (e.g. we can have 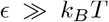 while 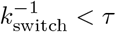). Since switching is independent of the 3-D conformation (and whether the TF is bound or not), it drives the system away from equilibrium, allowing behaviour that cannot be reproduced by a purely thermodynamic model (as discussed in the Discussion and Conclusions section, and see Ref. [24]).

### Simulation Details and Mapping to Real Units

We initialise the system from an ideal random walk for the chromatin polymer, and a random distribution of TFs. We evolve the system by performing Brownian Dynamics (BD) simulations where the solvent is implicitly modelled. Each particle (chromatin beads and transcription factors) obeys the following Langevin equation

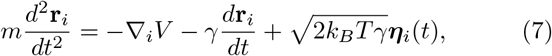

where r_*i*_ is the position of particle *i, γ* = *m/ξ* is the friction due to the solvent, where *ξ* is the velocity decorelation time. Here we take *ξ* = *τ*, where *τ* is the simulation time unit. Finally, *η_i_(t)* is a vector representing random uncorrelated noise, such that

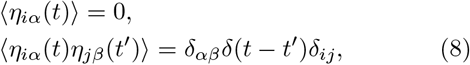

where *α* and *β* indicate Cartesian components, *δ_ij_* is Kro-necker’s delta, and *δ(t — t’)* denotes Dirac’s delta function. Eq. (7) is integrated with a constant time step Δ*t* = 0.01*τ*, for at least 70 × 10^6^ time steps while the system’s temperature is kept constant at a value *T* = 300 K. Eq. (7) is evolved using the Large-scale Atomic/Molecular Massively Parallel Simulator (LAMMPS) [60].

Loop extrusion and TF switching algorithms are implemented using in-house codes coupled to LAMMPS. Specifically, TF switching is performed by randomly selecting a fraction of the TFs and changing their state from “on” to “off” (or vice-versa), every 5*τ*.

In order to map simulation times to real units we match the diffusional properties of the chromatin fibre. The mean squared displacement (MSD) averaged over all simulated chromatin beads is measured, and we use this to fit the time unit to the experimentally measured MSD of fluorescently labelled chromatin loci in yeast cells [61]. The mapping varies slightly between the different models, but for simplicity we fix the time unit to *τ* = 50 ms in all cases.

Using this mapping, cohesin extrudes 2 beads (60 nm) in 250*τ* = 12.5 s, i.e. with a velocity of *v* ≃ 0.3 y*μ*m/min ≃ 30 kbp/min. TF switching occurs at rates ranging between 0 and 0.002 s^−1^.

### HiC data and boundary detection

HiC data for the GM12878 cell line were obtained at 10 kbp bin resolution, using “square root” normalization, from Ref. [9]. Simulated interaction maps were generated by recording contacts between chromatin beads whose 3D separation is < 5*σ* = 150 nm. These maps were then averaged over time (every 10^3^*τ* for the last 4 × 10^6^*τ*) and over 10 simulation replicas. The interaction values in the simulated maps range from 0 to 1, therefore when plotting simulated maps side-by-side with the HiC map, the later was rescaled by a factor of 400.

Boundaries were detected using an algorithm where the interactions map is analysed using a square window that slides along the diagonal and records the number of interactions inside the window as a function of its genomic position. Specifically, a 300 kbpx300 kbp window slides alongside the main diagonal with its corner positioned at the diagonal. This algorithm yields a profile of interactions with maxima at the center of domains and minima at the boundaries, hence a boundary is called for every local minima. The resulting set of boundaries is verified manually (via visual inspection) to correct wrong boundary calls due to the noisy signal of the interactions map. The same method was used to call boundaries in simulated contact maps. Boundary locations were said to have been correctly predicted if they appear in the simulated map within 54 kbp (6 beads) of the location on the HiC map. We used the Jaccard index, defined as

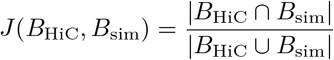

where *B*_HiC_ and *B*_sim_ are the sets of boundaries from the HiC data and simulations respectively. This takes values in the range from zero for no correctly predicted boundaries, to 1 for 100% agreement between simulation and HiC.

A quantification of the number of local and non-local interactions was performed by simply setting a threshold for locality and then calculating the total number of interactions recorded in the map between chromatin beads whose genomic separation is smaller (local interactions) or larger (non-local interactions) than the threshold. The ratio of the number of non-local to local interactions was then plotted as a function of the “locality” threshold. This ratio decreases as the threshold value increases.

Recent knock-out experiments for CTCF [34] and cohesin [25] were performed in liver and mouse embryonic stem cells respectively, so are not directly comparable to our human cell simulations. Nevertheless we show example interaction maps from this data in Fig. S3. We used these data sets to generate the plots in Fig. 7, showing the ratio of non-local to local interactions as a function of the threshold. As a suitable comparison for our simulations we selected the first 30 Mb section of mouse chromosome 7, which has a similar proportion of active/repressed and gene desert regions as our simulated section. We note that while the plots would differ for different chromosome sections, the trends between WT and knock-out are the same.

### “Virtual” 4C interaction profiles

“Virtual” 4C profiles were calculated for the HiC and simulation data using the “observed over expected” interaction maps (see caption of Fig. S2). Specifically, each diagonal line in the maps parallel to the main diagonal was first divided by the mean value of interactions in that diagonal. This way all interaction values have the same weight irrespective of their distance to the main diagonal. This facilitates the comparison between experimental and simulated profiles. Then, for the highlighted chromatin locus the number of interactions with all the other loci was calculated.

### From active to diffusive loop extruders

We compared two possible loop extrusion mechanisms: active extrusion, where the LE “heads” move apart unidirectionally at a specified speed, or diffusive extrusion where the two heads can move in either direction with equal probability. For the active case the extrusion speed is a parameter which can be chosen to give the best prediction of the data; for diffusing LEs, if we choose realistic parameters for diffusion, it takes much longer to generate the same size loops - we would need to run our simulations for infeasibly long times. In our previous work [29] (simulating much smaller chromatin segments) we showed that a simple 1-D model, where LE heads diffuse along a lattice, can accurately capture the behaviour of more detailed 3-D simulations with explicit diffusing slip-links.

#### 1-D simulations

Here we employ a similar 1-D model to the chromosome 7 segment. Chromatin is modelled as a 1-D lattice with N=10,000 lattice sites corresponding to the 10,000 simulated chromatin beads. The CTCF binding sites and motifs sequences used for the 3-D model are also used here, therefore we ran repeat simulations for the same stochastically chosen subsets of CTCF sites. Diffusing LEs are modelled as two heads that move independently, each occupying a lattice site. Every simulation step, each LE head moves to either neighbouring site with equal probability. Like in the 3-D model, LE heads cannot go past each other, or go past a CTCF oriented oppositely to their motion. Note that, within our rules, a LE head will go through a CTCF which is pointing away from it, but would subsequently halt if it diffuses back to the CTCF, as now this would be oriented towards the LE head. LEs (2 heads) attach randomly along the lattice at a rate *k*_on_ = 5 × 10^−5^ step^−1^, detach at a rate *k*_off_ = *k*_on_, and like in the 3-D case, if they form a CTCF loop (bring together two convergent CTCFs), they detach at a rate *k*_off_ = 0.1 × *k*_on_. The number of LEs is chosen so that the number of attached LEs is roughly 200 throughout the simulation. The 10 simulations ran for 20 × 10^6^ steps.

“HiC-like” interaction maps were calculated by considering that two lattice sites (chromatin loci) are in contact if they are occupied by matching LE heads. These maps were then averaged over time (every 400 steps) and over the 10 simulation replicas. Note that due to this definition of contacts it is not possible to obtain long-range interaction information.

#### 3-D simulations

We then performed 3-D simulations of diffusive loop extrusion, modelling the gene desert region chr7:10,000,000-20,000,000 at a lower resolution of 25 kbp per chromatin bead (N=400 beads). We derived the CTCF binding strengths according to the called ChIP-seq peak heights, and the CTCF relative orientations were assigned in the same way as described above. As before, all beads interact by a repulsive Weeks-Chandler-Andersen potential (Eq. 2), but here consecutive beads are bound together by a standard harmonic potential *U*_bond_(*r_ij_*) = (*K*_bond_/2)(*r_ij_* – *r*_0_)^2^, where we set *K*_bond_ = 1000k_*B*_*T* and an equilibrium distance 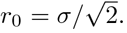. Additionally, all pairs of beads interact through an attractive Lennard-Jones potential (Eq. 5, with *ϵ* = *1k_B_T* and *r_thr_* = *2.5σ*), modelling the effect of macro-molecular crowding.

Like in the active extrusion case, diffusing LEs are modelled as harmonic bonds between two beads (Eq. 6), with *K*_LE_ = 10*k_B_T* and *r_0_* = *σ*. LEs are randomly positioned on the polymer, and diffusive extrusion is performed by allowing the two heads to slide independently along the polymer in both directions with equal probability (as in the 1-D model). LEs halt if they find a correctly oriented CTCF, with a probability set by the CTCF binding strength. Since a CTCF binding site can be visited several times for diffusive extrusion, we introduce a refractoriness time *τ*_reft_ = 10^2^*τ* for those CTCF sites where a possible bonding event fails. Once a LE halts at a CTCF, it remains stuck, meaning that the system will eventually “equilibrate” with “fixed” CTCF loops. In order to explore higher order loops, we introduce dissociation events where LEs unbind from CTCFs and start diffusing again (with *τ*_reft_ =4 × 10^3^*τ*). In the light of the active LE model, here LE-CTCF bonding events are such that only convergent CTCF loops are allowed.

The number of LEs in the simulation spans from 6 up to 15, and it is kept fixed during the simulation. To produce the initial polymer configurations, we generated random walk chains, where the distance between consecutive beads was fixed to 3*σ*, and let the system relax with a statistical minimization. The system obeys a Langevin equation (Eq. 7), which is integrated with a time-step Δ*t* = 0.005*τ*. An extrusion event occurs every 200Δ*t* = *τ*. We let the system evolve up to 4 × 10^4^*τ* and produce an ensemble of 10^2^ different conformations.

### Human chr4:20,300,000:22,600,000 simulations

We also analysed a different chromosome region - chr4:20,300,000:22,600,000, one of those thoroughly investigated in Ref. [20] - and compared the performance of the different loop extrusion models. We considered a polymer chain made of N=575 beads, each bead corresponding to 4 kbp, and we followed the same chromatin model and simulation details described in the previous section “3-D dLE simulations”. Here, we derived CTCF binding strengths and relative CTCF motif orientations using the standard approach described in Ref. [20].

First we implemented active Loop Extrusion (LE) with 3-D Molecular Dynamics (MD). Again we derived an ensemble of 10^2^ different replicates for the considered region and ran each simulation for 8 × 10^5^Δ*t*, corresponding to *t* = 4 × 10^3^*τ*. Finally, we computed the averaged contact maps over these configurations, and we considered two beads in contact if their 3-D separation was ≤ 1.5*σ*.

Second, we implemented the 3-D diffusive Loop Extrusion (dLE) model for this same region. We used a refractoriness time *τ*_reft_ = 10*τ* for the CTCF sites where a bonding event fails. Here, we derived an ensemble of 40 different configurations and we let the system evolve up to 10^6^Δ*t*.

And third, we investigated the same region with the “strings-and-binders” model [18]. We performed MD simulations considering a polymer chain made of N=2300 beads, each bead corresponding to 1 kbp. Here, all particles interact by a repulsive Weeks-Chandler-Andersen potential (Eq. 2) and consecutive beads are connected by a FENE spring (Eq. 1), while beads and binders interact by an attractive Lennard-Jones potential (Eq. 5). Chromatin is modelled by a homo-polymer, where all beads can interact with the same type of binders [62]. CTCF binding sites were considered according to the approach described in Ref. [20]. CTCF sites interact with an additional type of binders, which bridge CTCFs with opposite orientations (forward - reverse). For such a system, there are three possible thermodynamic states depending on the interaction energy and concentration of the binders - open, closed disordered, and closed ordered (for more details see Ref. [62]). As before, the system evolves under Langevin dynamics by MD with an integration timestep Δ*t* = 0.012*τ*. From the 3D equilibrium configurations in each thermodynamic state we computed averaged contact maps as described above (*r*_int_ = 3.5*σ*). Then, we find the mixture of the three states described above which best describes the locus by maximizing the distance corrected Spearman correlation coefficient between model and experimental data (at 4 kbp resolution, see also Refs. [51, 62]). We find the best mixture to be 10% open state and 90% closed state (of which 55% is in the ordered state and 35% in the disordered state).

**Figure S1:**
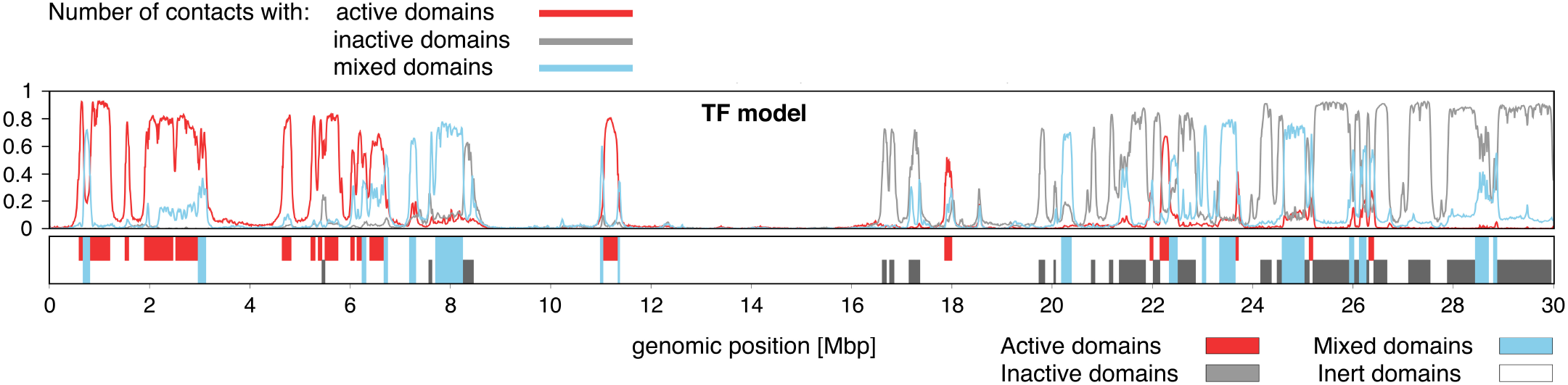
The TF model predicts A/B compartmentalisation. Plot showing, for each chromatin bead, the level of interaction with active (red), inactive (grey) and mixed (blue) domains. Chromatin domains interact mainly with other domains bearing the same histone marks, a typical feature of A/B compartmentalisation. For each domain the number of interactions with each type is normalised so that the total number of interactions sums to 1. The type of each domain is indicated below the plot. Domains were defined as the regions between domain boundaries, and their type was set according to the most frequent bead type in that region.

**Figure S2:**
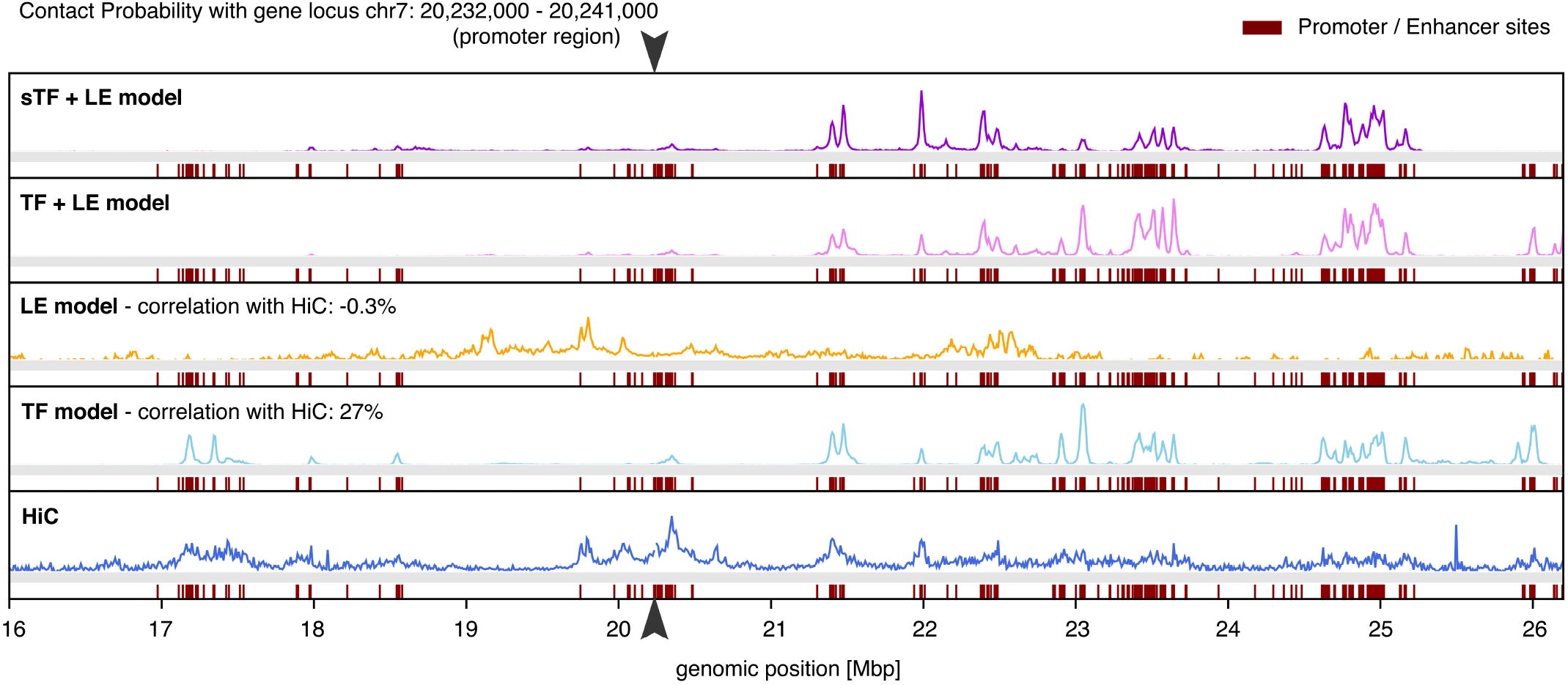
“Virtual” 4C of a locus for the various models. Plots showing interactions with a promoter at position chr7:20,232,000-20,241,000. The 4C-like interaction profiles were extracted from the “observed over expected” interactions map, where the expected value for a given pair of beads is the mean interaction level for all pairs of beads with the same genomic separation. Normalised in this way, it is easier to compare longer ranged interactions in the different models. The black arrowheads indicate the 4C viewpoint. Other enhancer/promoter sites are indicated in red below the curves.

**Figure S3:**
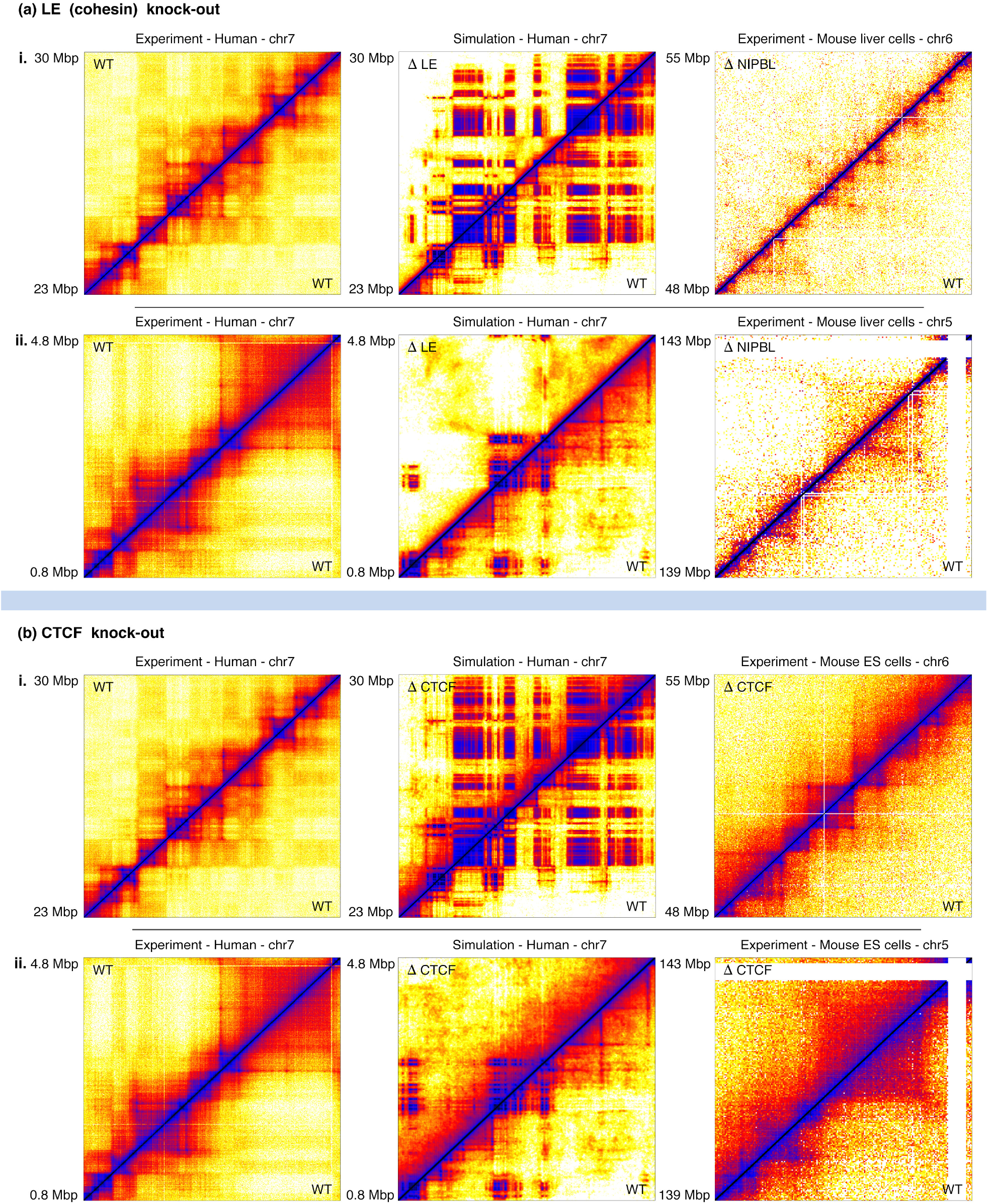
Comparison of knock-out experiments and simulations. (a) A cohesin knock-out is performed by removing LEs in simulations and the cohesin loader NIPBL in mouse liver cell experiments. Interaction maps are shown for two different chromosome regions - i and ii: (left) HiC map for a human chromosome 7 region, (centre) simulations’ map for the same human chromosome region, and (right) HiC map for a syntenic chromosome 6 region of mouse liver cells [25]. The human HiC map for the wild type is shown for reference. In the simulation and mouse experimental maps the knock-out (top left triangle) is compared with the wild type case (bottom right). (b) Similar interaction maps to the above, but for CTCF knock-out simulations (middle) and mouse embryonic stem cell experiments (right) [34].

**Figure S4:**
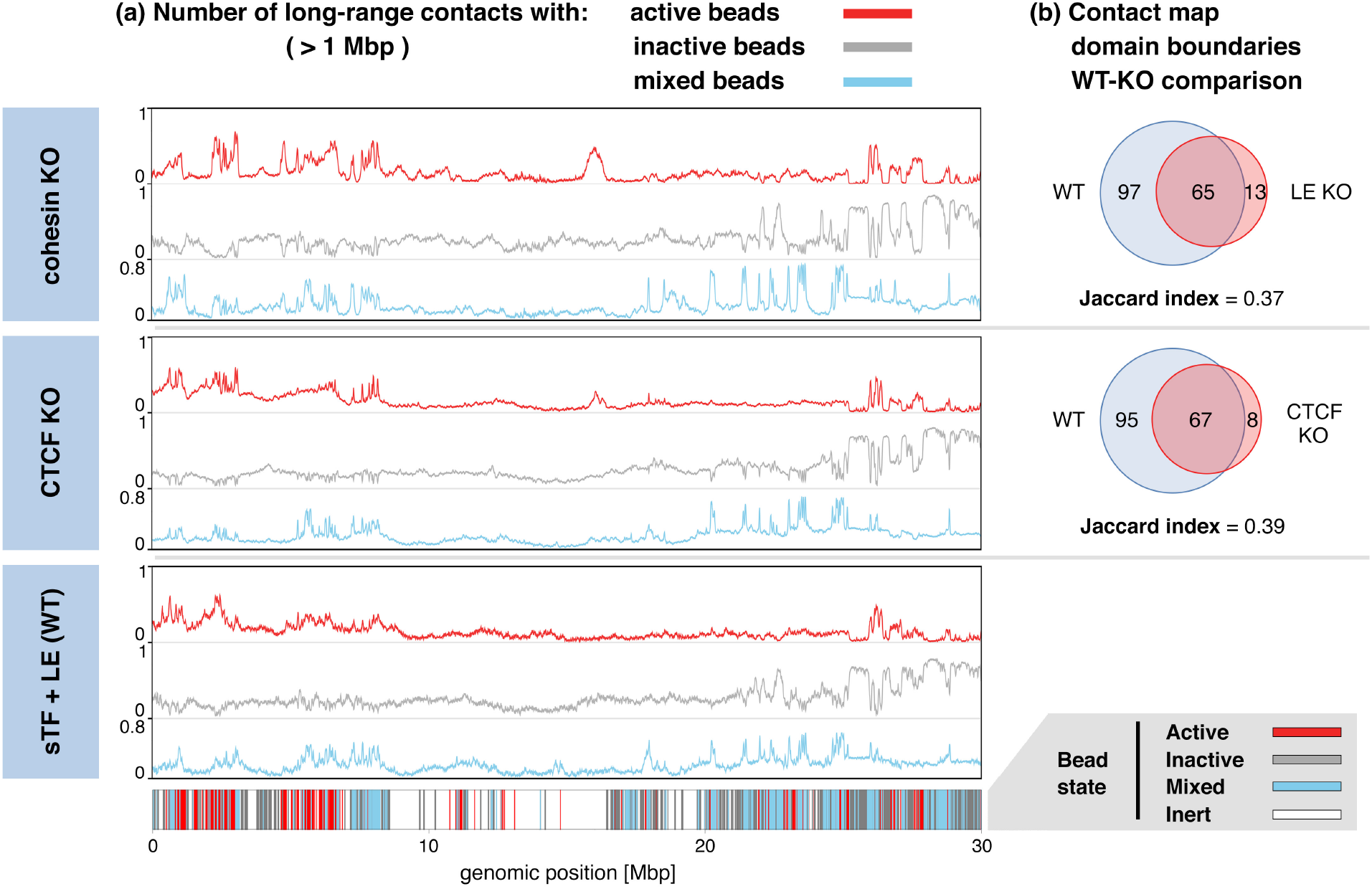
Quantitative comparison of knock-out and wild type simulations. As in Figure. 5: (a) Plots showing the level of interaction with active (red), inactive (grey) and mixed (blue) beads, for each chromatin bead. The labelling of each bead is indicated bellow the plots. (b) Venn diagrams showing the overlap between called domain boundaries in the KO and WT models.

**Figure S5:**
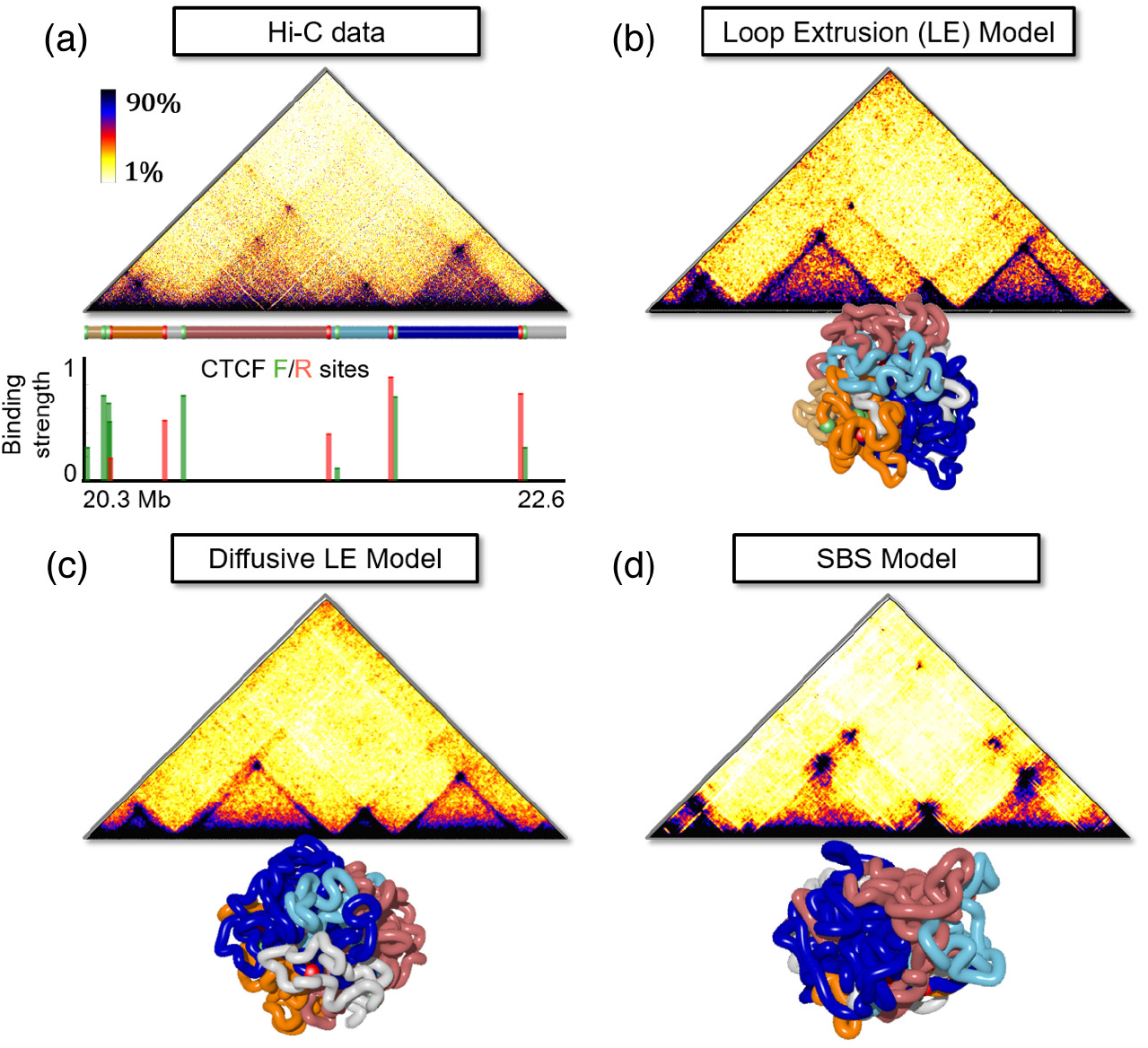
Performance of different polymer extrusion models to explain Hi-C data for the human chr4. (a) *in-situ* Hi-C data of the region chr4:20,300,000-22,600,000 for GM12878 cells at 4kb resolution investigated in Ref. [9] and corresponding CTCF binding sites, strength and orientation (green is forward, and red reverse). The coloured bar highlights CTCF positions and main polymer interacting regions to help 3-D visualization; Contact maps for the same region obtained with the (b) Loop Extrusion, (c) diffusive Loop Extrusion and (d) SBS 3-D chromatin models with interactions between CTCF sites oriented in opposite ways. For each model a typical 3D polymer structure is shown. Diffusive extrusion is as efficient as active extrusion at predicting HiC domain boundaries and peaks.

